# Mean-field modeling of brain-scale dynamics for the evaluation of EEG source-space networks

**DOI:** 10.1101/2020.09.16.299305

**Authors:** Sahar Allouch, Maxime Yochum, Aya Kabbara, Joan Duprez, Mohamad Khalil, Fabrice Wendling, Mahmoud Hassan, Julien Modolo

## Abstract

Understanding the dynamics of brain-scale functional networks at rest and during cognitive tasks is the subject of intense research efforts to unveil fundamental principles of brain functions. To estimate these large-scale brain networks, the emergent method called “electroencephalography (EEG) source connectivity” has generated increasing interest in the network neuroscience community, due to its ability to identify cortical brain networks with satisfactory spatio-temporal resolution, while reducing mixing and volume conduction effects. However, no consensus has been reached yet regarding a unified EEG source connectivity pipeline, and several methodological issues have to be carefully accounted for to avoid pitfalls. Thus, a validation toolbox that provides flexible “ground truth” models is needed for an objective methods/parameters evaluation and, thereby an optimization of the EEG source connectivity pipeline. In this paper, we show how a recently developed large-scale model of brain-scale activity, named COALIA, can provide to some extent such ground truth by providing realistic simulations of source-level and scalp-level activity. Using a bottom-up approach, the model bridges cortical micro-circuitry and large-scale network dynamics. Here, we provide an example of the potential use of COALIA to analyze, in the context of epileptiform activity, the effect of three key factors involved in the “EEG source connectivity” pipeline: (i) EEG sensors density, (ii) algorithm used to solve the inverse problem, and (iii) functional connectivity measure. Results showed that a high electrode density (at least 64 channels) is required to accurately estimate cortical networks. Regarding the inverse solution/connectivity measure combination, the best performance at high electrode density was obtained using the weighted minimum norm estimate (wMNE) combined with the weighted phase lag index (wPLI). Although those results are specific to the considered aforementioned context (epileptiform activity), we believe that this model-based approach can be successfully applied to other experimental questions/contexts. We aim at presenting a proof-of-concept of the interest of COALIA in the network neuroscience field, and its potential use in optimizing the EEG source-space network estimation pipeline.

## Introduction

There is now growing evidence suggesting that large-scale functional brain networks underlie complex brain functions during rest (Allen et al., 2014; Kabbara, Falou, Khalil, Wendling, & Hassan, 2017) and tasks (Hassan et al., 2015; O’Neill et al., 2017). Among the neuroimaging techniques used to derive the functional brain networks, the electroencephalography (EEG) technique provides a direct measure of electrical brain activity at the millisecond time scale. The past years have seen a noticeable increase of interest in “EEG source connectivity” methods to estimate brain networks at the cortical sources level while minimizing the volume conduction and field spread problems. Although consisting only of two main steps: 1) source reconstruction, and 2) connectivity assessment, there is still no consensus on a unified pipeline adapted to this approach, and many methodological questions remain unanswered. A first issue lies at the very first step of data recording with the question of optimal spatial resolution (i.e., density of sensors) needed to avoid misrepresentation of spatial information of brain activity (Song et al., 2015; Srinivasan, Tucker, & Murias, 1998). Another issue concerns the subsequent analysis: for each of the aforementioned steps, a large number of methods are available, each having its own properties, advantages and drawbacks, and addressing a different aspect of the data. An additional parameter warranting investigation is the spatial resolution of the reconstructed cortical sources (i.e., number of regions of interest) ranging from dozens to thousands of regions.

To tackle those challenges, several comparative studies have been conducted with the aim of evaluating the performance of the adopted techniques and the influence of different parameters affecting the network estimation procedure (Anzolin et al., 2019; Colclough et al., 2016; Fornito, Zalesky, & Bullmore, 2010; Halder, Talwar, Jaiswal, & Banerjee, 2019; Lantz, Grave de Peralta, Spinelli, Seeck, & Michel, 2003; Sohrabpour et al., 2015; Song et al., 2015; J. Wang et al., 2009).

In the context of EEG, several studies investigated the effect of different electrode montages on the estimation of functional connectivity. Increasing the number of electrodes has been shown to decrease the localization error in different contexts (Lantz et al., 2003; Sohrabpour et al., 2015; Song et al., 2015). (Song et al., 2015) recommended using 128 or 256 electrodes, while in (Sohrabpour et al., 2015) the most dramatic decrease in localization error was obtained when going from 32 to 64 electrodes. Other studies have focused on evaluating the performance of different inverse solutions using simulated and real EEG signals (Anzolin et al., 2019; Bradley, Yao, Dewald, & Richter, 2016; Grova et al., 2006; Halder et al., 2019). Compared methods include those based on the minimum norm estimate (MNE, LORETA, sLORETA, eLORETA, etc.) as well as beamformers (DICS, LCMV). However, to the best of our knowledge, there is no consensus yet on which inverse solution provides the most accurate results when estimating EEG-source-space networks. In the context of functional connectivity, the performance of various measures covering direct/indirect causal relations, marginal/partial associations, leakage correction, amplitude/phase coupling have been evaluated, and compared using either real data (Colclough et al., 2016), or simulated data in (H. E. Wang et al., 2014; Wendling, Ansari-Asl, Bartolomei, & Senhadji, 2009). Nevertheless, no agreement has been reached on which connectivity measure to adopt.

A challenging issue in such comparative studies resides in the absence of a ‘ground truth’ when dealing with real EEG data. Ideally, simultaneous scalp EEG and depth (intracranial) recordings are required, which is challenging to perform and is therefore unavailable in most studies. Thus, to overcome this issue, one possible solution is to use simulated data. It is worth mentioning that many studies have attempted to provide a ground truth for the validation of source reconstruction and connectivity estimation algorithms. For example, in (Schelter et al., 2006), a toy model was used in which the signal was considered as an oscillator and was driving the activity of other structures. However, such approach is limited in terms of spectral properties. A frequently adopted method is the use of multivariate autoregressive (MVAR) models as generator filters, in combination along with volume conductor head models to generate pseudo-EEG data (Anzolin et al., 2019; Haufe & Ewald, 2016). However, such models are linear and too simple compared to the complexity of actual brain activity. Another solution that overcomes some of the limitations of previous methods is the use of physiologically-inspired models. Here, we use a computational model named “COALIA” (Bensaid, Modolo, Merlet, Wendling, & Benquet, 2019), able to generate realistic brain-scale, cortical-level simulations while accounting for macro- (between regions) as well as the micro-circuitry (within a single region) of the human cortex, including the specificities of each neuronal type within each region. Scalp EEG signals can be then obtained through solving the EEG direct problem. We highlight the implications of this model in enhancing our interpretation of the reconstructed brain networks and in evaluating the key factors of the EEG source connectivity pipeline, such as 1) EEG sensor density, 2) solution of the EEG inverse problem, and 3) functional connectivity measure.

Here, we generate epileptiform, cortical activity (confined to the left hemisphere) and present a (not exhaustive) comparative study to evaluate, in this specific context, the effect of: 1) five different electrode densities (256, 128, 64, 32, 19); 2) two inverse solution algorithms, weighted minimum norm estimate (wMNE) and exact low resolution electromagnetic tomography (eLORETA); and 3) two functional connectivity measures, phase locking value (PLV) and weighted phase lag index (wPLI)) as they represent one of the most used combination of methods in the context of EEG source-space network estimation. We believe that the present study can be extended to address other methodological /experimental questions related to source connectivity estimation. We aim at presenting a proof-of-concept of the interest of COALIA in the network neuroscience field, and its potential use in optimizing the EEG source-space network estimation pipeline.

## Materials and Methods

The full pipeline of our study is summarized in Fig. 1.

**Fig 1.**
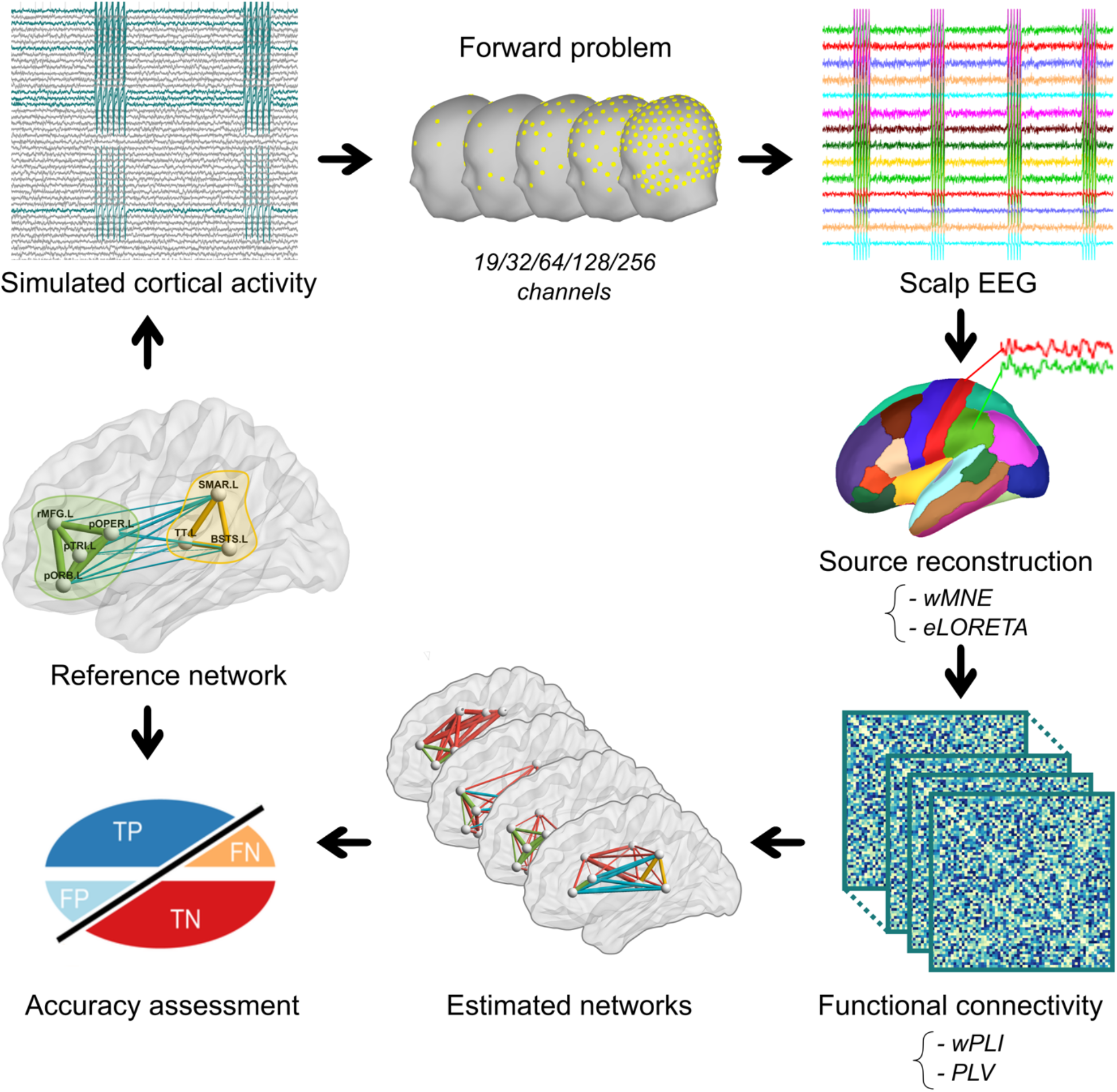
**Pipeline of the study. Cortical sources were simulated using COALIA. The forward model was solved for five electrode montages (19, 32, 64, 128, 256 electrodes). Scalp EEG signals were generated. Cortical sources were reconstructed using wMNE and eLORETA as inverse solutions. Functional connectivity between reconstructed sources was assessed over 30 trials using PLV and wPLI algorithms. Accuracy was computed to assess the performance of the network estimation.**

### Simulations

Source-space brain activity was generated using a physiologically-grounded computational model, named COALIA. As aforementioned, it generates brain-scale electrophysiology activity while accounting for the macro-(between regions) as well as the micro-circuitry (within a single region) of the brain (for details, see (Bensaid et al., 2019)). We considered a scenario inspired from a general scheme of the organization of human partial seizures presented in (Bartolomei, Guye, & Wendling, 2013), and proposing the existence of an epileptogenic subnetwork as well as a propagation subnetwork. In the present study, the two subnetworks were located in the left hemisphere. The epileptogenic subnetwork included four cortical regions: the rostral middle frontal gyrus, pars opercularis, pars triangularis, and pars orbitalis; and the propagation subnetwork included the supramarginal, banks superior temporal sulcus, and transverse temporal cortex. Regions affiliations were based on the Desikan-Killiany atlas (Desikan et al., 2006). Epileptiform activity was generated in the epileptogenic and propagation subnetworks, while background activity was assigned to the remaining cortical regions. A detailed description of the model along with along with all simulation parameters relative to the data are provided in the Supplementary Materials and the generated source signals are provided in the GitHub repository. All sources belonging to a single patch were synchronized at a zero lag, while a delay of 30 ms was introduced between the two subnetworks to reflect the propagation of spikes between relatively distant regions in the brain. A timeseries of ~6 min at 2048 Hz was simulated, and was segmented into 10-second epochs. A total of 30 epochs was selected for the subsequent analysis. This value was chosen based on a previous study (Hassan et al., 2017), dealing with similar issues based on computational modeling and epileptic spikes.

### EEG Electrodes density and direct problem

Five different electrode montages were used to generate scalp EEG signals. We selected the GSN HydroCel EEG configuration (EGI, Electrical geodesic Inc) for the 256, 128, 64 and 32 channels density, as well as the international 10-20 system (Klem, Lüders, Jasper, & Elger, 1999) for the 19 channels array. For each electrode configuration, the lead field matrix describing the electrical and geometrical characteristics of the head was computed for a realistic head model using the Boundary Element Method (BEM) using OpenMEEG (Gramfort, Papadopoulo, Olivi, & Clerc, 2010) implemented in the Brainstorm toolbox (Tadel, Baillet, Mosher, Pantazis, & Leahy, 2011) for Matlab (The Mathworks, USA, version 2018b). To generate EEG scalp simulations, we solved the forward problem as follows:

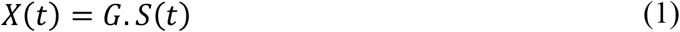

where *X*(*t*), *S*(*t*) are scalp EEG and cortical timeseries respectively, and *G* the lead field matrix. We used only the lead field vectors reflecting the contribution of the sources located at the centroid of the regions of interest defined on the basis of the Desikan-Killiany atlas (Desikan et al., 2006) (right and left insula were excluded, leaving 66 regions of interest).

Finally, in order to simulate measurement noise, spatially and temporally uncorrelated signals were added to the scalp EEG as follows (Anzolin et al., 2019):

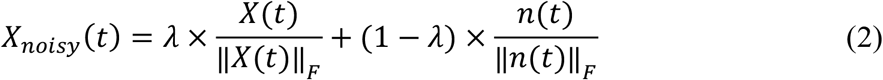

where *X*(*t*) are the scalp EEG and *n*(*t*) is the white uncorrelated noise. ∥*X*(*t*)∥_*F*_ and ∥*n* (*t*)∥_*F*_ refers to the Frobenius norm of the multivariate time series *X*(*t*) and *n*(*t*) respectively. First, *λ* was fixed to 1 (i.e. no measurement noise was added). Second, for evaluating the different methods in the presence of noise, *λ* was varied between 0.85 and 0.95 with a 0.01 step.

### EEG inverse problem

Solving the EEG inverse problem consists of estimating the position, orientation and magnitude of dipolar sources *Ŝ*(*t*). Cortical sources were positioned at the centroids of Desikan-killiany regions, and oriented normally to the cortical surface. Thus, the inverse problem was reduced to the computation of the magnitude of dipolar sources *Ŝ*(*t*):

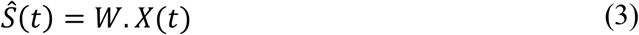

where *X*(*t*) is the scalp EEG. Several algorithms have been proposed to solve this problem and estimate W based on different assumptions related to the spatiotemporal properties of the sources and regularization constraints (see (Baillet, Mosher, & Leahy, 2001) for a review). Here, we used two methods widely used in EEG source connectivity analysis: the weighted minimum norm estimate (wMNE) and the exact low-resolution electromagnetic tomography (eLORETA).

### Weighted Minimum Norm Estimate (wMNE)

The minimum norm estimate (MNE) originally proposed by (Hämäläinen & Ilmoniemi, 1994)h searches for a solution that fits the measurements with a least square error. The wMNE (Fuchs, Wagner, Köhler, & Wischmann, 1999; Lin et al., 2006) compensates for the tendency of MNE to favor weak and surface sources:

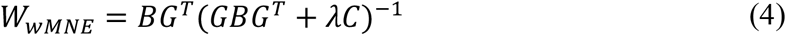

where *λ* is the regularization parameter and *C* is the noise covariance matrix computed, in our case, from the pre-spikes baseline. The matrix *B* is a diagonal matrix built from matrix G with non-zero terms inversely proportional to the norm of lead field vectors. It adjusts the properties of the solution by reducing the bias inherent to the standard MNE solution:

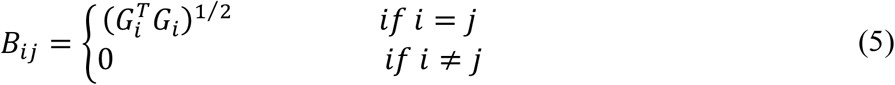

### Exact low-resolution brain electromagnetic tomography (eLORETA)

The exact low-resolution electromagnetic tomography (eLORETA) belongs to the family of weighted minimum norm inverse solutions. However, it does not only account for depth bias, it also has exact zero error localization in the presence of measurement and structured biological noise (Pascual-Marqui, 2007):

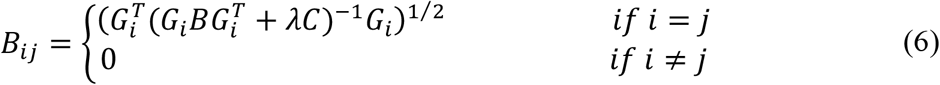

eLORETA was originally described using the whole brain volume as source space. However, in the present study, in order to facilitate the comparison with other methods, we restricted the source space to the cortical surface. Regarding the regularization parameters of wMNE and eLORETA, we used default values included in Brainstorm and fieldtrip toolboxes.

### Connectivity measures

We evaluated in this study two of the most popular connectivity metrics, both based on the assessment of the phase synchrony between regional time-courses, namely PLV (phase-locking value) and PLI (phase-lag index), as detailed below.

### Phase-locking value

For two signals *x*(*t*) and *y*(*t*), the phase-locking value (Lachaux et al., 2000) is defined as:

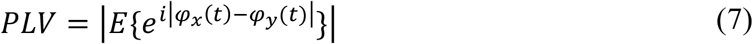

where *E*{.} is the expected value operator and *φ*(*t*) is the instantaneous phase derived from the Hilbert transform.

### Weighted phase-lag index

While the phase-lag index (PLI) quantifies the asymmetry of the phase difference, rendering it insensitive to shared signals at zero phase lag (Stam, Nolte, & Daffertshofer, 2007) that supposedly induce spurious volume conduction effects, the weighted PLI (wPLI) attempts to further weight the metric away from zero-lag contributions (Vinck, Oostenveld, Van Wingerden, Battaglia, & Pennartz, 2011).

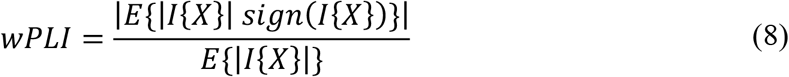

where *I*{*X*} denotes the imaginary part of the signal’s cross-spectrum.

Connectivity matrices were computed in broadband [1-45 Hz] for all considered electrode densities and possible inverse solution/connectivity combinations, resulting in 20 connectivity matrices for each epoch. The resulting matrices were thresholded by keeping nodes with the highest 12% strength values, corresponding to the proportion of nodes originally used to simulate the 2 subnetworks. A node’s strength was defined as the sum of the weights of its corresponding edges.

### Results quantification

In order to assess the performance of each investigated parameter (i.e., electrodes number, inverse solution, connectivity measure), the accuracy of the estimated networks with respect to the ground truth was computed as follows:

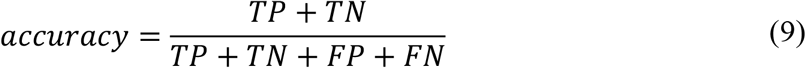

where *TP* (i.e., true positive) represents the connections present in the reference as well as in the estimated network, *TN* (i.e., true negative) refers to the absent connections in both the reference and estimated networks, *FP* (i.e., false positive) represents the connections obtained in the estimated network exclusively, and *FN* stands for the links missing in the estimated network. Accuracy values range between 0 and 1.

### Statistical analysis

Statistical analyses were performed using R (R Core Team, 2020). We used linear mixed model analyses to investigate the effects of electrode number, inverse solution method, and connectivity measure on the accuracy of the estimated networks. Mixed models have several advantages, such as the ability to account for the dependence between the different measures, and to model random effects (see (Gueorguieva & Krystal, 2004)). We used the *lmer* function of the {*lme4*} package (Bates, Mächler, Bolker, & Walker, 2015) with the following model that includes, epoch, electrode number, inverse solution method and connectivity measures as interacting fixed effects, and also a random intercept for epochs:

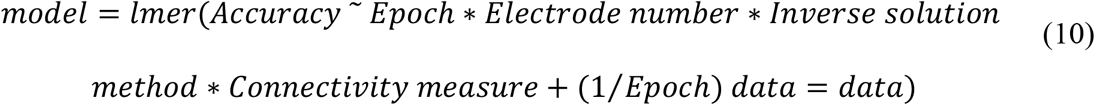

We applied a square root transform to the data, since this led to a better compliance of the model with the assumptions of normality and homoscedasticity of model’s residuals than for raw data. Calculation of the significance of the fixed effects was performed using the anova function of the {*car}* package that computes ANOVA F-tests (Fox & Weisberg, 2019). In order to assess the quality of the model, we computed marginal and conditional R^2^ that were obtained from the {*MuMin*}package. In case of significant main effects, we performed *post-hoc* analyses using the *glht* function of the {*multcomp*} package that calculates adjusted p-values using individual z tests (Hothorn, Bretz, & Westfall, 2008). The significance threshold was set to *p* = 0.05.

## Results

For each sensor density and inverse solution connectivity measure, estimated networks averaged over trials are illustrated in Fig. 2. Those results illustrate clearly that the accuracy of the estimated networks was dramatically influenced by scalp sensors density. The higher the number of electrodes, the more accurate the reconstructed networks were. Also, the inverse solution/connectivity measures combinations performed distinctively. The best performance was obtained using wMNE/wPLI at 64, 128, and 256 electrodes. wMNE/PLV and eLORETA/PLV performed similarly and were slightly less accurate than wMNE/wPLI at a high sensor density. However, eLORETA/wPLI exhibited the least estimation accuracy even with a high number of electrodes. The accuracy values for all electrodes montage and inverse solution/connectivity measures combinations were plotted in Fig. 3. The influence of the sensor density was confirmed by the statistical analysis with a significant sensor density effect (*F*_(4,532)_ = 333.53, *p* < 0.001, *conditional R*^2^ = 0.84, *marginal R*^2^ = 0.82). *Post-hoc* analyses showed significant accuracy improvement when using 256 electrodes as compared to 64 (*p* < 0.05), 32 (*p* < 0.001), and 19 (*p* < 0.01) electrodes. Increasing the sensor density from 128 electrodes to 256 electrodes did not provide further benefit.

**Fig 2.**
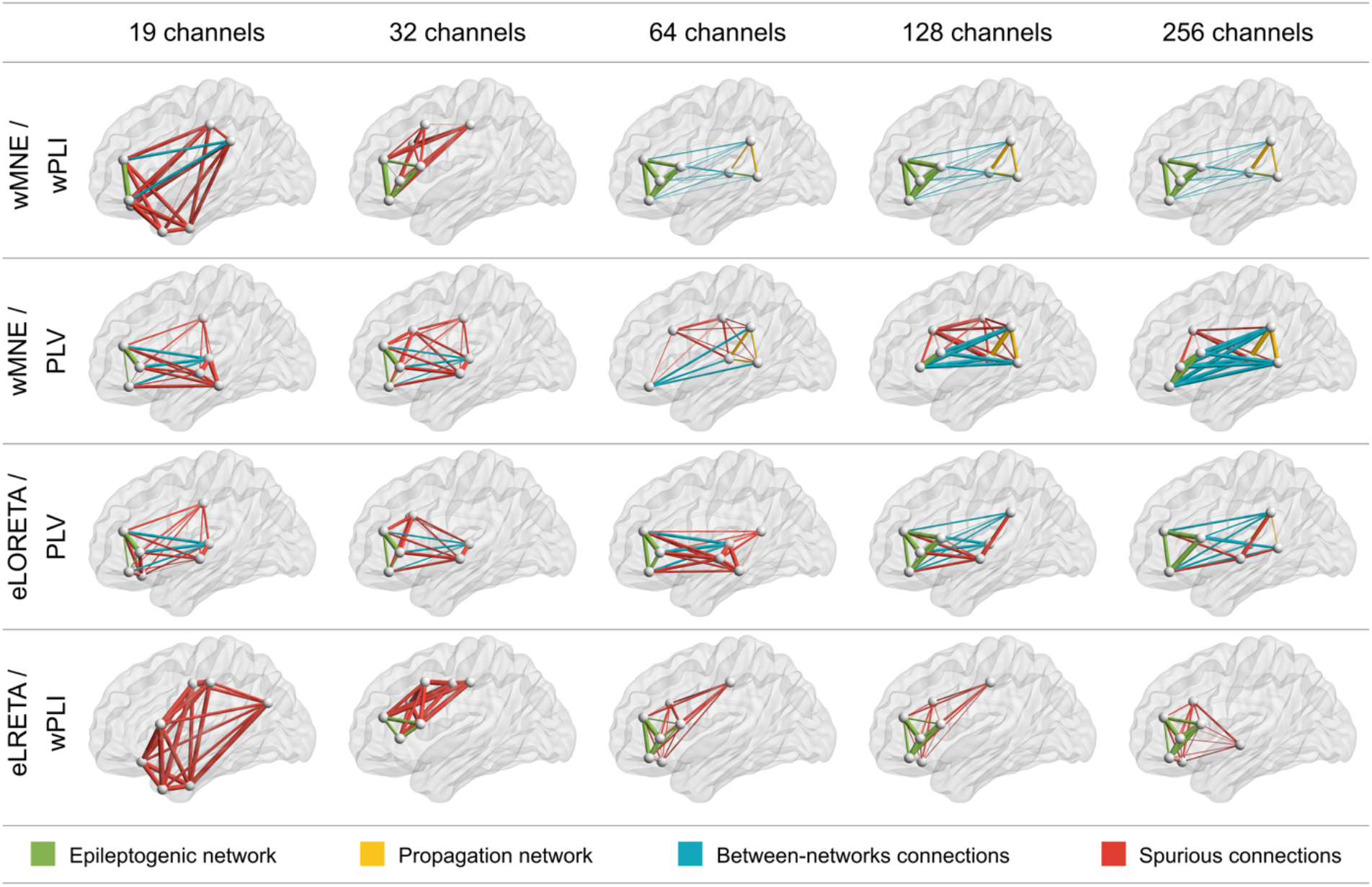
**Average networks over trials for all electrode montages and inverse solution/connectivity measure combinations. Networks were thresholded by keeping nodes with the highest 12% strength values, which corresponds to the proportion of nodes originally present in the reference network. Connections in green and yellow belong to the epileptogenic and propagation subnetworks respectively. Connections in blue represent the connectivity between the two subnetworks. Connections in red are spurious connections, that do not exist in the reference network.**

**Fig 3.**
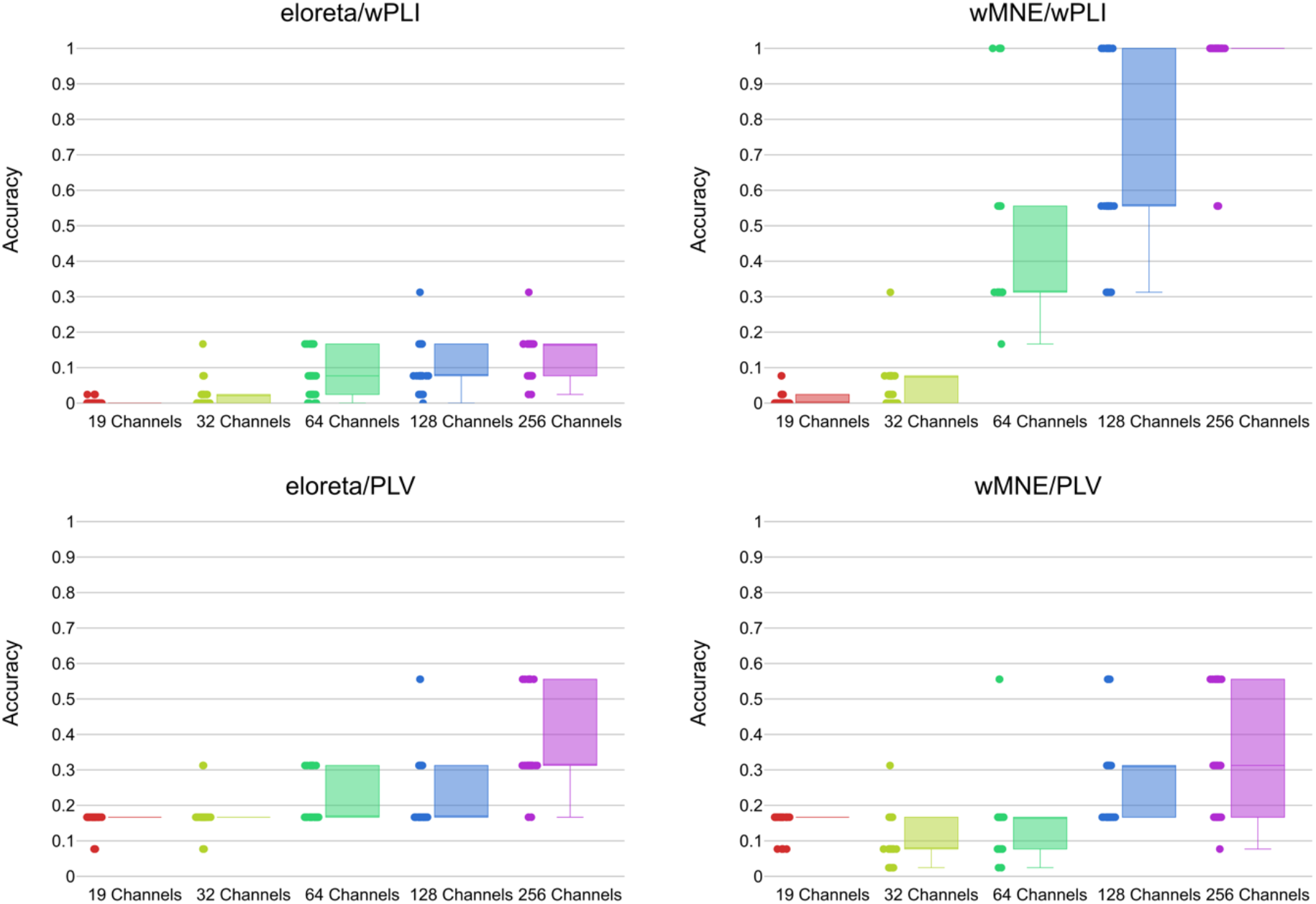
**Accuracy of the estimated networks based on different electrode montages for each inverse solution/connectivity measure.**

A non-significant difference was obtained in all other cases. Differences between results obtained with 19 electrodes and those obtained with 32, 64, or 128 electrodes were all non-significant. Similarly, no differences were detected between 32 and 64, 32 and 128, 64 and 128 electrode montages. Regarding the inverse solution, wMNE significantly outperformed eLORETA (*F*_(1,532)_ = 281.75, *p* < 0.001, *conditional R*^2^ = 0.84, *marginal R*^2^ = 0.82). Also, statistical analyses showed a significant effect of the connectivity measure (*F*_(1,532)_ = 83.19, *p* < 0.001, *conditional R*^2^ = 0.84, *marginal R*^2^ = 0.82). The accuracy of the estimated networks was slightly higher with wPLI than with PLV. Interestingly, the combination inverse solution/connectivity measure combination had also a significant effect on the network estimation accuracy (*F*_(1,532)_ = 478.91, *p* < 0.001, *conditional R*^2^ = 0.84, *marginal R*^2^ = 0.82).

The highest network estimation accuracy was reached using wMNE/wPLI, while the worst performance was obtained with eLORETA/wPLI. eLORETA/PLV and wMNE/PLV had similar average accuracy values. *Post-hoc* analyses showed significant difference between wMNE/wPLI and both eLORETA/PLV (*p* < 0.001) and wMNE/PLV (*p* < 0.001). Similarly, results obtained with eLORETA/wPLI were significantly different from those obtained with eLORETA/PLV (*p* < 0.001) and wMNE/PLV (*p* < 0.001). On the other hand, differences between eLORERA/PLV and wMNE/PLV and between eLORETA/wPLI and wMNE/wPLI were not statistically significant. All the detailed results of the statistical analysis can be found in the Supplementary Materials.

In Fig. 4, the mean accuracy and standard error of each inverse solution/connectivity measure combination are plotted against different levels of noise (see Materials and Methods) for the case of 256 electrodes. With the exception of wMNE/PLV, the different combination methods maintained a relatively stable performance at different levels of noise. Plots relative to other electrode montages were also included in the Supplementary Materials.

**Fig 4.**
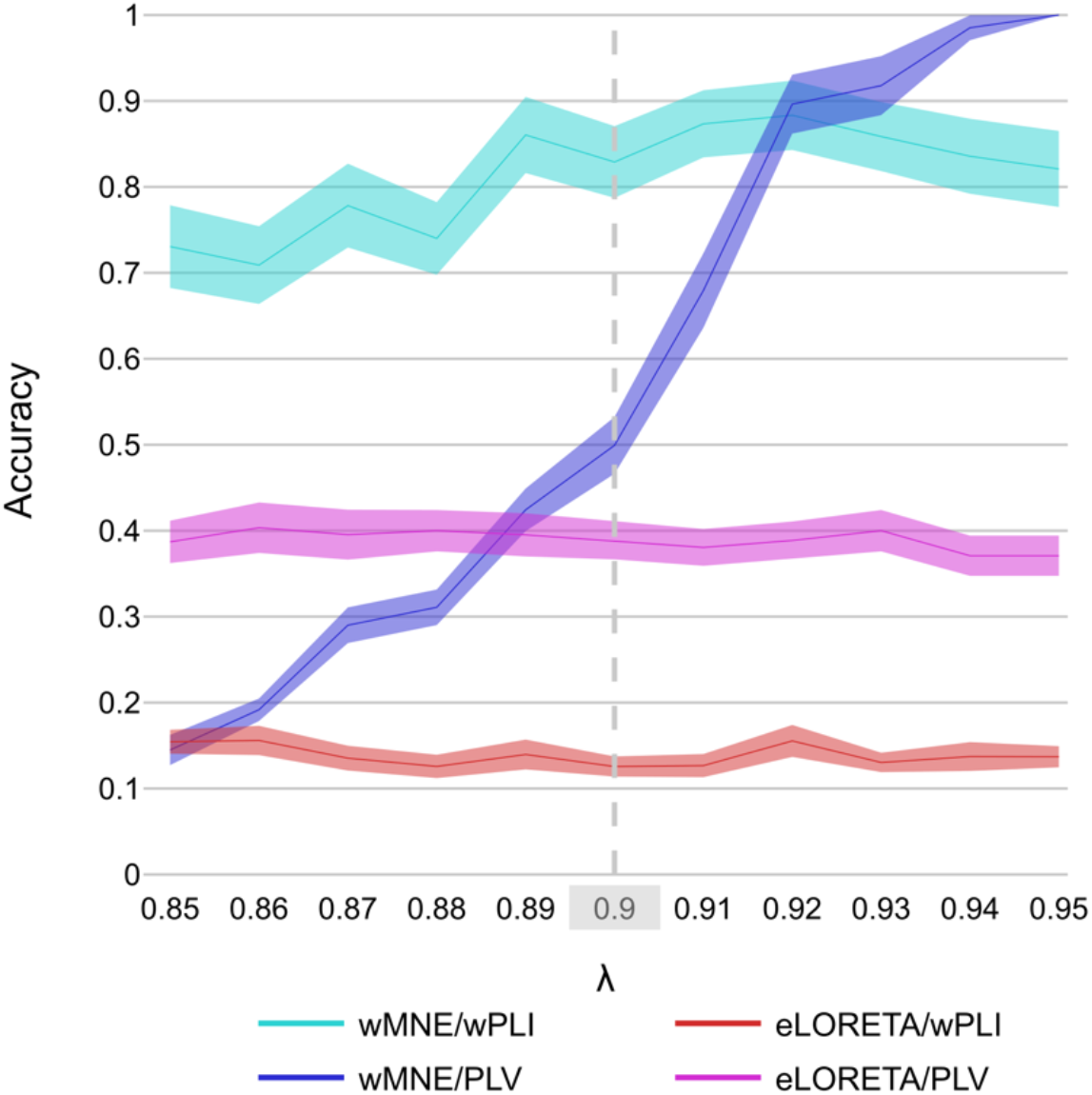
**Mean accuracy and standard error of each inverse solution/connectivity measure combination plotted against different levels noise for the case of 256 electrodes**

## Discussion

To the best of our knowledge, there is still no consensus on the most optimized pipeline for reconstructing EEG source-space networks. At each step of this pipeline, several methods have indeed been proposed and many parameters need to be defined. Several comparative studies have investigated different methods/parameters affecting the estimation of functional networks. A key challenge in such studies (i.e., when dealing with real EEG data) is the difficulty to obtain a ground truth, which prevents the exact evaluation of the performance of each considered method. In order to overcome this issue, in this paper we propose to use a recently developed, physiologically-grounded computational model, and highlight its potential use in optimizing the EEG network estimation procedure. Our objective was to provide a proof-of-concept regarding the use of COALIA for testing methods and parameters included in the source connectivity estimation. We believe that it can be used to address many of the methodological considerations related to EEG source connectivity estimation, and ultimately provide a validation of the pipeline, a step of increased interest, especially with the emergence of the relatively new field called “network neuroscience” (Bassett & Sporns, 2017). In this study, we used the model to simulate cortical-level sources from which scalp-EEG signals were generated, and then evaluated the effects of EEG channels density, two source reconstruction algorithms, and connectivity measures. Specifically, as a first step of this strategy to provide a ground truth using COALIA, we considered a scenario consisting of an epileptogenic and propagation network where epileptiform activity is present.

Overall, results obtained for the five considered electrode montages demonstrate clearly that the spatial resolution of the sensor array dramatically affects the accuracy of network estimation: as expected, increasing spatial resolution involves a higher accuracy of reconstructed networks. These results were expected theoretically, and are in line with previous studies (Lantz et al., 2003; Sohrabpour et al., 2015; Song et al., 2015). Recording EEG data with a low sensor density array can indeed contribute to a misrepresentation of high spatial frequency signal as a low spatial frequency signal. Therefore, to avoid aliasing the Nyquist criteria (*F*_*S*_, > 2 ∗ *F*_*max*_) should be respected and high spatial resolution is required (i.e., small interelectrode distance) (Song et al., 2015; Srinivasan et al., 1998). Interestingly, increasing the number of electrodes from 128 to 256 did not provide a significant improvement. On a different note, (Song et al., 2015) found that adding sensors on the inferior surface of the head (including the neck and the face) improves localization accuracy, even with sparse arrays. Therefore, it may be interesting to study the effect of the head coverage provided by different sensor array layouts, and not only the number of electrodes in each array.

Comparing inverse solutions and connectivity measures showed that wMNE performed better than eLORETA, and wPLI performed better than PLV. In contrast to our results, in (Tait, Szul, & Zhang, 2020), eLORETA outperformed wMNE at both voxel and ROI level. Even though (Colclough et al., 2016) did not recommend phase-based metrics as a first choice for assessing MEG functional connectivity, they were in favor of using measures that are not affected by zero-lag phase coupling. Interestingly, our study showed that a more crucial parameter is the combination of inverse method and connectivity measure. Although wMNE/wPLI performed better than eLORETA/wPLI, the PLV connectivity measure performed similarly with both eLORETA and wMNE methods. Thus, the choice of the inverse solution and connectivity measure is recommended to be made simultaneously. In a previous study (Hassan, Dufor, Merlet, Berrou, & Wendling, 2014), wMNE/PLV combination had the best performance in the context of a picture naming task. This combination has also showed better performance than other combinations (eLORETA and wPLI were not included) when applied to simulated epileptic spikes (Hassan et al., 2017). A possible difference between the current simulations and (Hassan et al., 2017) is that, in the latter, the reference network were very dense locally with a very high number of zero-lag correlations which may favor methods that do not remove these connections (such as PLV). Moreover, neither eLORETA nor wPLI were investigated in that study. It is therefore worth noting that the results obtained in this study are specific to the analyzed condition, i.e., epileptic spikes: therefore, we cannot be certain that the same combination of methods will provide the best network estimation accuracy when analyzing networks related to cognitive tasks or resting state, such as the alpha/beta DMN for instance (i.e., physiological activity), which is the main objective of the future steps of this work.

Finally, we varied *λ* between 0.85 and 0.95 to evaluate the effect of different noise levels on the performance of the inverse solutions and connectivity estimates. Our results have showed that the additive scalp-level noise mostly affected wMNE/PLV, while other methods had a more stable performance. In (Anzolin et al., 2019; Haufe & Ewald, 2016), *λ* was fixed at 0.9. Results reported in Fig. 4 show that, at this specific value, wMNE/PLV reached a good performance (mean = 0.50 as compared to wMNE/wPLI (mean = 0.83), eLORETA/PLV (mean = 0.39), and eLORETA/wPLI (mean = 0.13). One can also notice that at higher SNR (*λ* > 0.9), wMNE/PLV outperforms other combinations. These observations highlight the importance of applying an effective preprocessing of EEG signals before reconstructing the cortical networks, as well as a robust source connectivity method to correctly estimate functional connectivity. A more detailed assessment of the effect of noise levels on the performance of source connectivity algorithms is a possible future avenue of research.

## Methodological considerations

Here, our objective was to provide a typical example of the use of the COALIA model to investigate the effect of different pipeline-related parameters on EEG source-space network analysis. With our approach, we aimed at promoting the use of computational modeling as a ground-truth to evaluate parameters of EEG source connectivity methods. Using this approach, other parameters could be also evaluated and other scenarios could be also generated, and we suggest hereafter possible extensions for this work. First, we simulated in this study a network with 7 regions generating spike activity, while background brain activity was attributed to all other regions. However, it would be even more realistic for the network neuroscience field to use the model to simulate different rhythms of resting-state data (alpha/beta-band activity in the DMN network, for instance) and then evaluate the desired techniques in such context, rather than restricting the study to spikes/background activity scenarios. Second, the inverse solutions compared in this paper both belong to the family of minimum norm estimates methods. Other algorithms based on beamformers, such as the widely used linearly constrained minimum variance (LCMV) were not tested here. Moreover, it is worthy to mention that the two connectivity measures included in this study estimate phase synchrony between regional time-series. Other existing methods investigate instead the amplitude correlation between signals, such as the amplitude envelope correlation (AEC), which is widely used in the context of MEG functional connectivity. In a large study investigating the reliability of different connectivity metrics (Colclough et al., 2016), Colclough et al. have suggested that AEC between orthogonalized signals is the most consistent connectivity measure to employ in the context of resting-state recordings. Thus, it is noteworthy that the results obtained here do not necessarily extend to other inverse solutions or connectivity measures, nor they are generalized to all experimental context. Moreover, the connectivity assessed here using PLV and wPLI is bidirectional, however, since we introduced a time delay between the two subnetworks, we propose that studying directional connectivity metrics (e.g., Granger causality) may also lead to additional insights specifically related to Bartolomei’s model, where epileptic activity is transferred from the epileptogenic network to the propagation network.

In order to threshold connectivity matrices, only the nodes with the highest 12% strength were kept. This proportional threshold was chosen to ensure that the number of nodes in the estimated networks matches the number of nodes within the reference network (7/66=12%). Obviously, this choice is not completely realistic, since we cannot have any *a priori* in terms of experimental data on the exact number of activated brain regions. However, adopting a proportional, rather than a statistical thresholding for example, ensures that the density of estimated networks matches that of the ground truth, which is necessary for the correct assessment of the accuracy of estimated networks in our case (van den Heuvel et al., 2017).

Let us mention that we used in this study the accuracy to quantify the difference between estimated and reference networks. Other network-based metrics can be also useful to compute the similarities between these networks (Mheich et al., 2018; Mheich, Wendling, & Hassan, 2020). Also, the present study is limited to cortical regions based on the assumption that sub-cortical regions are not easily accessible from scalp EEG recordings. However, it has been proved that the performance of some inverse algorithms and connectivity estimators depends on the position of the reconstructed sources (Anzolin et al., 2019). Thus, a more extensive study comparing source connectivity approaches should include the effect of the location of sources.

In terms of head model, we have built for the purpose of this study a realistic head model for the ICBM152 MRI template that consisted in three nested homogeneous mesh surfaces shaping the brain (642 vertices), skull (642 vertices) and scalp (1,082 vertices) with conductivity values of 0.33 Sm^−1^, 0.0042 Sm^−1^ and 0.33 Sm^−1^, respectively. However, source connectivity analysis in EEG/MEG are usually affected by the choice of the head model describing electrical and geometrical characteristics of the head (simplified/realistic head models, individual/template MRI, tissue types, tissue conductivity) (Cho, Vorwerk, Wolters, & Knösche, 2015; Wolters et al., 2006). Thus, it may be worthy to use COALIA and benefit from the presence of a ground truth to examine the influence of the head model on EEG source connectivity analysis. Finally, functional connectivity was estimated, here, in broadband [1 – 45] Hz. However, it can be also computed in each of the classical EEG frequency sub-bands (i.e., delta, theta, alpha, beta, gamma) separately (Bettus et al., 2008; Canuet et al., 2011) which could be done in a more exhaustive study.

## Conclusion

In this work, we have provided evidence that COALIA, a recently developed, physiologically-inspired computational model can provide a ground-truth for comparative studies aiming at optimizing the EEG-source connectivity pipeline. Using this model-based approach, several methodological questions can be addressed. Here, we assessed the effect of the number of EEG electrodes, as well as the inverse solution/connectivity measure combination in the context of simulated epileptic activity. Our results suggest that a higher network estimation accuracy requires a high number of EEG electrodes, and suggest a careful choice of an efficient inverse solution/connectivity measure combination.

## Funding

This work was financed by Rennes University, the Institute of Clinical Neurosciences of Rennes (project named EEGCog). It was also supported by the Programme Hubert Curien CEDRE (PROJECT No. 42257YA), the National Council for Scientific Research (CNRS-L) and the Agence Universitaire de la Francophonie (AUF) and the Lebanese university.

## Conflicts of interest/Competing interest

The authors have no conflicts of interest to declare that are relevant to the content of this article.

## Availability of data and material

Data used in this work can be found at https://github.com/sahar-allouch/comp-epi.git.

## Code availability

Data and Codes supporting the results of this study are available at https://github.com/sahar-allouch/comp-epi.git. We used Matlab (The Mathworks, USA, version 2018b), Brainstorm toolbox (Tadel et al., 2011), Fieldtrip toolbox ((Oostenveld, Fries, Maris, & Schoffelen, 2011); http://fieldtriptoolbox.org), OpenMEEG (Gramfort et al., 2010) implemented in Brainstorm, and BrainNet Viewer (Xia, Wang, & He, 2013).

## Acknowledgments

This work was financed by the Rennes University, the Institute of Clinical Neuroscience of Rennes (project named EEGCog). Authors would also like to thank the Lebanese Association for Scientific Research (LASER) and Campus France, Programme Hubert Curien CEDRE (PROJECT No. 42257YA), for supporting this study. The authors would like to acknowledge the Lebanese National Council for Scientific Research (CNRS-L), the Agence Universitaire de la Francophonie (AUF) and the Lebanese university for granting Ms. Allouch a doctoral scholarship.

## Notes

### Competing Interest Statement

The authors have declared no competing interest.

## References

Allen, E. A., Damaraju, E., Plis, S. M., Erhardt, E. B., Eichele, T., & Calhoun, V. D. (2014). Tracking Whole-Brain Connectivity Dynamics in the Resting State. Cerebral Cortex, 24, 663–676. https://doi.org/10.1093/cercor/bhs352

Anzolin, A., Presti, P., Van De Steen, F., Astolfi, L., Haufe, S., & Marinazzo, D. (2019). Quantifying the Effect of Demixing Approaches on Directed Connectivity Estimated Between Reconstructed EEG Sources. Brain Topography. https://doi.org/10.1007/s10548-019-00705-z

Baillet, S., Mosher, J. C., & Leahy, R. M. (2001). Electromagnetic Brain Mapping. IEEE Signal Processing Magazine, (November), 14–30.

Bartolomei, F., Guye, M., & Wendling, F. (2013). Abnormal binding and disruption in large scale networks involved in human partial seizures. EPJ Nonlinear Biomedical Physics, 1(4), 1–16. https://doi.org/10.1140/epjnbp11

Bassett, D. S., & Sporns, O. (2017). Network neuroscience. Nature Neuroscience, 20(3), 353–364. https://doi.org/10.1038/nn.4502

Bates, D., Mächler, M., Bolker, B. M., & Walker, S. C. (2015). Fitting linear mixed-effects models using lme4. Journal of Statistical Software, 67(1). https://doi.org/10.18637/jss.v067.i01

Bensaid, S., Modolo, J., Merlet, I., Wendling, F., & Benquet, P. (2019). COALIA: A Computational Model of Human EEG for Consciousness Research. Frontiers in Systems Neuroscience, 13, 1–18. https://doi.org/10.3389/fnsys.2019.00059

Bettus, G., Wendling, F., Guye, M., Valton, L., Régis, J., Chauvel, P., & Bartolomei, F. (2008). Enhanced EEG functional connectivity in mesial temporal lobe epilepsy. Epilepsy Research, 81(1), 58–68. https://doi.org/10.1016/j.eplepsyres.2008.04.020

Bradley, A., Yao, J., Dewald, J., & Richter, C. P. (2016). Evaluation of electroencephalography source localization algorithms with multiple cortical sources. PLoS ONE, 11(1), 1–14. https://doi.org/10.1371/journal.pone.0147266

Canuet, L., Ishii, R., Pascual-Marqui, R. D., Iwase, M., Kurimoto, R., Aoki, Y., … Takeda, M. (2011). Resting-state EEG source localization and functional connectivity in schizophrenia-like psychosis of epilepsy. PLoS ONE, 6(11). https://doi.org/10.1371/journal.pone.0027863

Cho, J. H., Vorwerk, J., Wolters, C. H., & Knösche, T. R. (2015). Influence of the head model on EEG and MEG source connectivity analyses. NeuroImage, 110, 60–77. https://doi.org/10.1016/j.neuroimage.2015.01.043

Colclough, G. L., Woolrich, M. W., Tewarie, P. K., Brookes, M. J., Quinn, A. J., & Smith, S. M. (2016). How reliable are MEG resting-state connectivity metrics? NeuroImage, 138, 284–293. https://doi.org/10.1016/j.neuroimage.2016.05.070

Desikan, R. S., Ségonne, F., Fischl, B., Quinn, B. T., Dickerson, B. C., Blacker, D., … Killiany, R. J. (2006). An automated labeling system for subdividing the human cerebral cortex on MRI scans into gyral based regions of interest. NeuroImage, 31, 968–980. https://doi.org/10.1016/j.neuroimage.2006.01.021

Fornito, A., Zalesky, A., & Bullmore, E. T. (2010). Network scaling effects in graph analytic studies of human resting-state fMRI data. Frontiers in Systems Neuroscience, 4, 1–16. https://doi.org/10.3389/fnsys.2010.00022

Fox, J., & Weisberg, S. (2019). An R Companion to Applied Regression (Third). Sage. https://doi.org/10.1177/0049124105277200

Fuchs, M., Wagner, M., Köhler, T., & Wischmann, H. A. (1999). Linear and nonlinear current density reconstructions. Journal of Clinical Neurophysiology, 16(3), 267–295. https://doi.org/10.1097/00004691-199905000-00006

Gramfort, A., Papadopoulo, T., Olivi, E., & Clerc, M. (2010). OpenMEEG: opensource software for quasistatic bioelectromagnetics. BioMedical Engineering OnLine, 9(45). https://doi.org/10.1186/1475-925X-8-1

Grova, C., Daunizeau, J., Lina, J. M., Bénar, C. G., Benali, H., & Gotman, J. (2006). Evaluation of EEG localization methods using realistic simulations of interictal spikes. NeuroImage, 29, 734–753. https://doi.org/10.1016/j.neuroimage.2005.08.053

Gueorguieva, R., & Krystal, J. H. (2004). Move over ANOVA? Progress in Analyzing Repeated-Measures Data and Its Reflection in Papers Published in the Archives of General Psychiatry. Arch Gen Psychiatry, 61, 310–317. Retrieved from http://odur.let.rug.nl/~nerbonne/teach/rema-stats-meth-seminar/presentations/Strik-AIC-2011-May-17.pdf

Halder, T., Talwar, S., Jaiswal, A. K., & Banerjee, A. (2019). Quantitative evaluation in estimating sources underlying brain oscillations using current source density methods and beamformer approaches. ENeuro, 6(4), 1–14. https://doi.org/10.1523/ENEURO.0170-19.2019

Hämäläinen, M. S., & Ilmoniemi, R. J. (1994). Inetrpreting magnetic fields of the brain: minimum norm estimates. Medical & Biological Engineering & Computing, 32, 35–42.

Hassan, M., Benquet, P., Biraben, A., Berrou, C., Dufor, O., & Wendling, F. (2015). Dynamic reorganization of functional brain networks during picture naming. Cortex, 73, 276–288. https://doi.org/10.1016/j.cortex.2015.08.019

Hassan, M., Dufor, O., Merlet, I., Berrou, C., & Wendling, F. (2014). EEG source connectivity analysis: From dense array recordings to brain networks. PLoS ONE, 9(8). https://doi.org/10.1371/journal.pone.0105041

Hassan, M., Merlet, I., Mheich, A., Kabbara, A., Biraben, A., Nica, A., & Wendling, F. (2017). Identification of Interictal Epileptic Networks from Dense-EEG. Brain Topography, 30(1), 60–76. https://doi.org/10.1007/s10548-016-0517-z

Haufe, S., & Ewald, A. (2016). A Simulation Framework for Benchmarking EEG-Based Brain Connectivity Estimation Methodologies. Brain Topography, 32(4), 625–642. https://doi.org/10.1007/s10548-016-0498-y

Hothorn, T., Bretz, F., & Westfall, P. (2008). Simultaneous inference in general parametric models. Biometrical Journal, 50(3), 346–363. https://doi.org/10.1002/bimj.200810425

Kabbara, A., Falou, W. E. L., Khalil, M., Wendling, F., & Hassan, M. (2017). The dynamic functional core network of the human brain at rest. (August 2016), 1–16. https://doi.org/10.1038/s41598-017-03420-6

Klem, G. H., Lüders, H. O., Jasper, H. H., & Elger, C. (1999). The ten-twenty electrode system of the International Federation. The International Federation of Clinical Neurophysiology. Electroencephalogr. Clin. Neurophysiol. Suppl.

Lachaux, J.-P., Rodriguez, E., Le Van Quyen, M., Lutz, A., Martinerie, J., & Varela, F. J. (2000). Studying Single-Trials of Phase Synchronous Activity in the Brain. International Journal of Bifurcation and Chaos, 10(10), 2429–2439. https://doi.org/10.1142/s0218127400001560

Lantz, G., Grave de Peralta, R., Spinelli, L., Seeck, M., & Michel, C. M. (2003). Epileptic source localization with high density EEG: How many electrodes are needed? Clinical Neurophysiology, 114(1), 63–69. https://doi.org/10.1016/S1388-2457(02)00337-1

Lin, F. H., Witzel, T., Ahlfors, S. P., Stufflebeam, S. M., Belliveau, J. W., & Hämäläinen, M. S. (2006). Assessing and improving the spatial accuracy in MEG source localization by depth-weighted minimum-norm estimates. NeuroImage, 31, 160–171. https://doi.org/10.1016/j.neuroimage.2005.11.054

Mheich, A., Hassan, M., Khalil, M., Gripon, V., Dufor, O., & Wendling, F. (2018). SimiNet: A Novel Method for Quantifying Brain Network Similarity. IEEE Transactions on Pattern Analysis and Machine Intelligence, 40(9), 2238–2249. https://doi.org/10.1109/TPAMI.2017.2750160

Mheich, A., Wendling, F., & Hassan, M. (2020). Brain network similarity: Methods and applications. Network Neuroscience, 4(3), 507–527. https://doi.org/10.1162/netn_a_00133

O’Neill, G. C., Tewarie, P. K., Colclough, G. L., Gascoyne, L. E., Hunt, B. A. E., Morris, P. G., … Brookes, M. J. (2017). Measurement of dynamic task related functional networks using MEG. NeuroImage, 146, 667–678. https://doi.org/10.1016/j.neuroimage.2016.08.061

Oostenveld, R., Fries, P., Maris, E., & Schoffelen, J. M. (2011). FieldTrip: Open source software for advanced analysis of MEG, EEG, and invasive electrophysiological data. Computational Intelligence and Neuroscience, 2011. https://doi.org/10.1155/2011/156869

Pascual-Marqui, R. D. (2007). Discrete, 3D distributed, linear imaging methods of electric neuronal activity. Part 1: exact, zero error localization. Retrieved from http://arxiv.org/abs/0710.3341

Schelter, B., Winterhalder, M., Hellwig, B., Guschlbauer, B., Lücking, C. H., & Timmer, J. (2006). Direct or indirect? Graphical models for neural oscillators. Journal of Physiology Paris, 99(1), 37–46. https://doi.org/10.1016/j.jphysparis.2005.06.006

Sohrabpour, A., Lu, Y., Kankirawatana, P., Blount, J., Kim, H., & He, B. (2015). Effect of EEG electrode number on epileptic source localization in pediatric patients. Clinical Neurophysiology, 126, 472– 480. https://doi.org/10.1016/j.clinph.2014.05.038

Song, J., Davey, C., Poulsen, C., Luu, P., Turovets, S., Anderson, E., … Tucker, D. (2015). EEG source localization: Sensor density and head surface coverage. Journal of Neuroscience Methods, 256, 9–21. https://doi.org/10.1016/j.jneumeth.2015.08.015

Srinivasan, R., Tucker, D. M., & Murias, M. (1998). Estimating the spatial Nyquist of the human EEG. Behavior Research Methods, Instruments, and Computers, 30(1), 8–19. https://doi.org/10.3758/BF03209412

Stam, C. J., Nolte, G., & Daffertshofer, A. (2007). Phase lag index: Assessment of functional connectivity from multi channel EEG and MEG with diminished bias from common sources. Human Brain Mapping, 28, 1178–1193. https://doi.org/10.1002/hbm.20346

Tadel, F., Baillet, S., Mosher, J. C., Pantazis, D., & Leahy, R. M. (2011). Brainstorm: A user-friendly application for MEG/EEG analysis. Computational Intelligence and Neuroscience, 2011. https://doi.org/10.1155/2011/879716

Tait, L., Szul, M. J., & Zhang, J. (2020). Cortical source imaging of resting-state MEG with a high resolution atlas : An evaluation of methods. https://doi.org/10.1101/2020.01.12.903302

van den Heuvel, M. P., de Lange, S. C., Zalesky, A., Seguin, C., Yeo, B. T. T., & Schmidt, R. (2017). Proportional thresholding in resting-state fMRI functional connectivity networks and consequences for patient-control connectome studies: Issues and recommendations. NeuroImage, 152(February), 437–449. https://doi.org/10.1016/j.neuroimage.2017.02.005

Vinck, M., Oostenveld, R., Van Wingerden, M., Battaglia, F., & Pennartz, C. M. A. (2011). An improved index of phase-synchronization for electrophysiological data in the presence of volume-conduction, noise and sample-size bias. NeuroImage, 55(4), 1548–1565. https://doi.org/10.1016/j.neuroimage.2011.01.055

Wang, H. E., Bénar, C. G., Quilichini, P. P., Friston, K. J., Jirsa, V. K., & Bernard, C. (2014). A systematic framework for functional connectivity measures. Frontiers in Neuroscience, 8. https://doi.org/10.3389/fnins.2014.00405

Wang, J., Wang, L., Zang, Y., Yang, H., Tang, H., Gong, Q., … He, Y. (2009). Parcellation-dependent small-world brain functional networks: A resting-state fmri study. Human Brain Mapping, 30, 1511–1523. https://doi.org/10.1002/hbm.20623

Wendling, F., Ansari-Asl, K., Bartolomei, F., & Senhadji, L. (2009). From EEG signals to brain connectivity: A model-based evaluation of interdependence measures. Journal of Neuroscience Methods, 183, 9–18. https://doi.org/10.1016/j.jneumeth.2009.04.021

Wolters, C. H., Anwander, A., Tricoche, X., Weinstein, D., Koch, M. A., & MacLeod, R. S. (2006). Influence of tissue conductivity anisotropy on EEG/MEG field and return current computation in a realistic head model: A simulation and visualization study using high-resolution finite element modeling. NeuroImage, 30(3), 813–826. https://doi.org/10.1016/j.neuroimage.2005.10.014

Xia, M., Wang, J., & He, Y. (2013). BrainNet Viewer: A Network Visualization Tool for Human Brain Connectomics. PLoS ONE, 8(7). https://doi.org/10.1371/journal.pone.0068910

